# Extensive patient-to-patient single nuclei transcriptome heterogeneity in pheochromocytomas and paragangliomas

**DOI:** 10.1101/2022.05.05.489848

**Authors:** Peter Brazda, Cristian Ruiz-Moreno, Wout Megchelenbrink, Henri J L M Timmers, Hendrik G. Stunnenberg

## Abstract

Pheochromocytomas (PC) and paragangliomas (PG) are rare neuroendocrine tumors of varied genetic makeup, associated with high cardiovascular morbidity and a variable risk of malignancy. The source of the transcriptional heterogeneity of the disease and the underlying biological processes determining the outcome in PCPG remains largely unclear. We focused on PCPG tumors with germline SDHB and RET mutations, representing distinct prognostic groups with worse or better prognoses, respectively. We applied single-nuclei RNA sequencing (snRNA-seq) on tissue samples from 11 patients and found high patient-to-patient transcriptome heterogeneity of the neuroendocrine tumor cells. The tumor microenvironment also showed heterogeneous profiles mainly contributed by macrophages of the immune cell clusters and Schwann cells of the stroma. Performing non-negative matrix factorization we identified common transcriptional programs active in RET and SDHB as well as distinct modules including neuronal development, hormone synthesis and secretion, and DNA replication. Comparison of the SDHB and RET transcriptomes with that of developmental stages of adrenal gland formation suggests different developmental stages at which PC and PG tumors appear to be arrested.

## INTRODUCTION

Pheochromocytomas (PC) and sympathetic paragangliomas (PG) are rare neuroendocrine tumors, originating from chromaffin cell-related populations located inside or outside the adrenal glands, respectively. PCPG is associated with significant morbidity and mortality [1]. The current therapy of choice is surgical resection; however, the disease can be associated with a lifelong risk of tumor persistence or recurrence [2].

A plethora of genes has been reported to be responsible for a diverse hereditary background in up to 40% of PCPG [3, 4]. Based on the bulk transcriptional and genomic profiles, PCPG has been divided into two major classes. Tumors in class 1 are predominantly extra-adrenal and display germline mutations in the succinate dehydrogenase complex (SDHB, SDHC, SDHD collectively referred to as SDHx), the most common form of PCPG. SDHx tumors have the worst prognosis with a 30–70% risk of metastasis or recurrence [5]. Class 2 PCPG detected in 5% of hereditary PCPGs comprise amongst others germline and/or somatic mutations of the RET proto-oncogene and have a better prognosis.

In this study, we exploited recent advances in single-nuclei RNA-seq to compare the gene expression landscapes of PCPG with SDHB and RET germline mutations and explore the transcriptional heterogeneity and to gain insight into the molecular basis of their different prognosis.

## MATERIALS AND METHODS

### Preparation of Single-Nuclei Suspensions

Previously selected tissue blocks were transferred for the RadboudUMC biobank and stored at −80°C. Nuclei were prepared from frozen tissue under RNAse-free conditions. Briefly, samples were cut to ~7 mm pieces, while kept on dry-ice. The pieces were minced in a pre-cooled douncer in 500uL ice-cold Nuclei EZ Lysis buffer 5x with pestle-A and 10x with pestle-B. The suspension was passed through a 70 μm cell strainer and washed with 1.5 mL cold Nuclei EZ Lysis and incubated on ice for 5’. The lysate was washed in Nuclei wash/resuspension buffer (1xPBS completed with 1% BSA and 0.2U/ul RNAsin Plus (Promega, #N2611) and passed through a 40 μm cell strainer. Nuclei were stained with DAPI. To exclude doublets and debris from the final mix and to precisely determine the number of loaded nuclei, we applied FACS. 15000 nuclei were sorted into a pre-cooled tube containing the RT-mix (RT-reagent + TSO + Reducing agent B), right before loading the mix to one lane of the Chromium chip, 8.3 ul RT-enzyme was added to the mix, according to the standard protocol of the Chromium Single Cell 3’ kit (v2). All the following steps for the library preparation were performed according to the manufacturer’s protocol. Paired-end sequencing was used to sequence the prepared libraries using an Illumina NextSeq sequencer.

### Single-Cell RNA-seq Data Processing and Quality Control (QC)

Raw sequencing data were converted to FASTQ files with bcl2fastq. Reads were aligned to the human genome reference sequence (GRCH38) and counted with STAR. The CellRanger (10X Genomics) analysis pipeline was used to sample demultiplexing and single cell gene counting to generate the gene-cell expression matrix for each library. The gene expression matrix was then processed and analyzed by *Seurat* package in R. To filter out low-quality cells, we first removed cells (nuclei) for which less than 300 or more than 4000 genes were detected. The cell count and gene count information for single cell datasets of the PCPG samples are listed in Table1.

### Dimensionality Reduction, Clustering and Visualization

Data were normalized by sequencing depth, scaled to 10000 counts, log-transformed, and regressed against the UMI-counts and percentage using the ScaleData function of the Seurat package. Principal components analysis was performed on the scaled data with the 4000 most variable genes. Using 15 first principal components, we calculated a UMAP representation of the data for visualization and calculated clusters using the *FindNeighbors* and *FindClusters* functions with the resolution parameter set to 0.3. Marker genes differentiating between the clusters were identified with the FindAllMarkers function. Before running a second round of clustering after sub-setting the original dataset to a cell type of interest, we applied the DietSeurat function to collect the unmodified expression matrix of the subset of cells, without any transformations.

To identity cell types, we used sets of well-established marker and annotated each cell type based on their average expression. Cluster (or clusters) marker genes were determined with the *FindAllMarkers* function and required to be expressed in at least 25% of the cells in a cluster with a minimal log expression difference of 0.25 between clusters.

### Inferred CNV Analysis from snRNA-seq

Large-scale copy number variations (CNVs) were inferred from single-nuclei gene expression profiles using the *inferCNV* package [6] using the i3 HMM parameter, a window size of 101 genes and the “cluster_by_groups” parameter is true. To identify the distinct chromosomal gene expression pattern of neuroendocrine cells, all other cell were set as the “reference” cells. CNVs in the reference cells would still be detectable.

### Expression Programs of Intra-tumoral Heterogeneity

We applied non-negative matrix factorization (NMF) via the *RunNMF* function of the swne [7] package to extract transcriptional programs of malignant cells of each sample. We set the number of factors to 28 for each sample. For each of the resulting factors, we considered the top 50 genes with the highest NMF scores as characteristics of that given factor. We used the *AddModuleScore* function in the Seurat package to evaluate the degree to which individual cells express a certain pre-defined expression program, and thus determine the scores. All tumor cells were then scored according to the 280 NMF programs. Hierarchical clustering of the scores for each program using Pearson correlation coefficients as the distance metric and Ward’s linkage revealed 10 correlated sets of metaprograms. The gene list of 10 meta-programs can be found in TableS5.

### Logistic Regression for Similarity Calculation

To measure the similarity of a target single-cell transcriptome to a reference single-cell dataset, we used the logistic regression method described in [8]. Briefly, we trained a logistic regression model with elastic net regularization (α = 0.6) on the reference training set. We then used this trained model to infer a similarity score for each cell in the query dataset for each cell type in the reference data. Predicted logits were averaged within each cluster or sample group in the query dataset.

## RESULTS

We performed single-nuclei transcriptomic profiling (snRNA-seq) on resected tumor tissues from 11 treatment-naïve patients to generate a comprehensive PCPG atlas. Molecular diagnoses revealed germline RET and SDHB mutations in 5 and 6 patients, respectively (**Table S1**). All RET-PCPG samples were retrieved from the adrenal gland, while the SDHB-PCPG tumors are from various locations, including the bladder, the adrenal gland, the retroperitoneal- and the mediastinal area.

### Cell Type Composition of PCPG Tissue

Stringent quality filtering yielded 50,868 nuclei with an average of 1,800 genes detected per nuclei (**Methods**, **Table S1**). The merged expression profiles were compressed into a 2D-coordinate system using Uniform Manifold Approximation and Projection for Dimension Reduction (UMAP). The cells were grouped into 20 clusters and were annotated by their location, mutation group as well as patient ID **(Fig1A, Table S2)**.

**Figure1.**
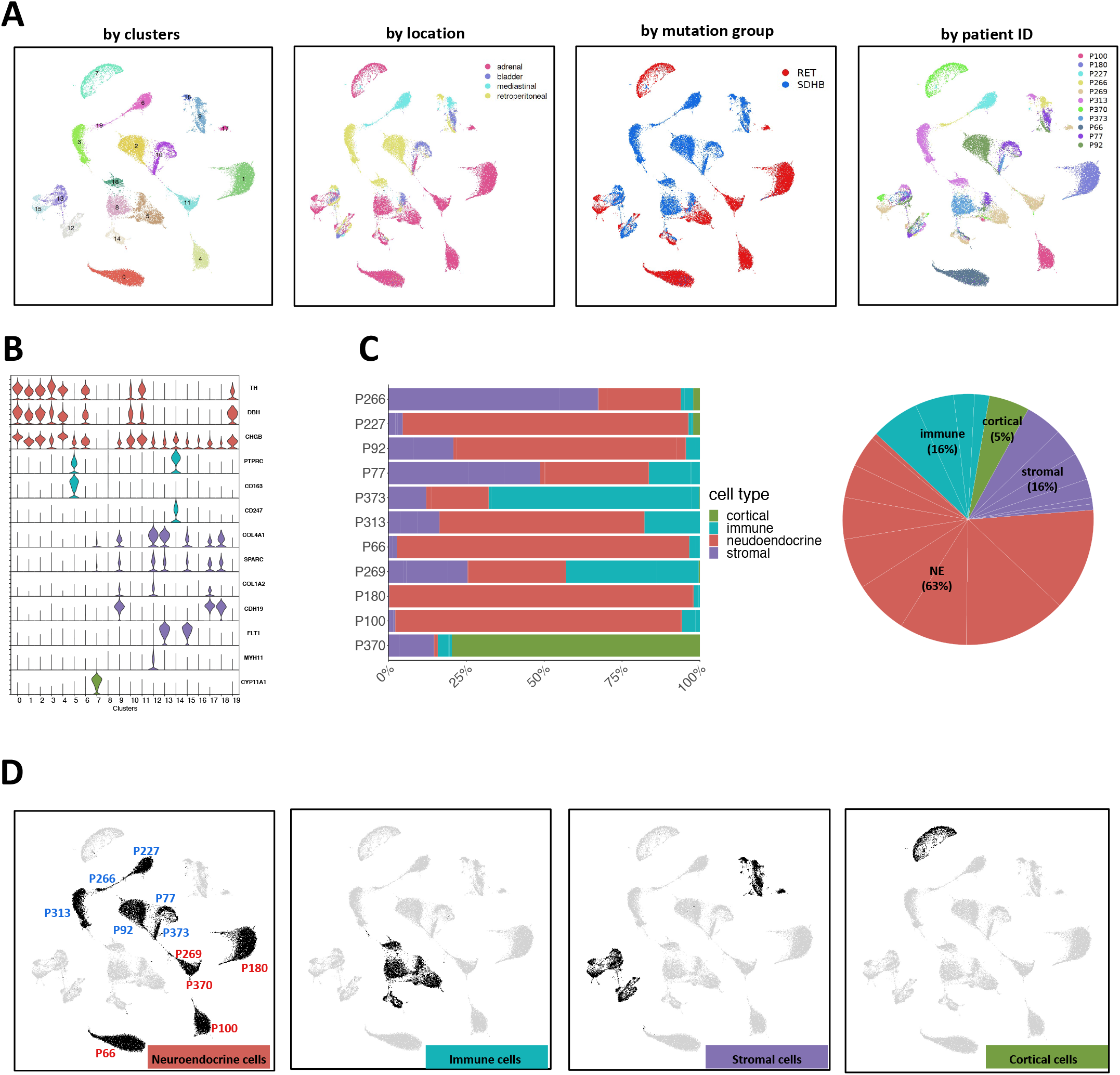
A. UMAP visualization of all the 50,868 cells grouped according to their cluster annotation and colored by their clusters, location of origin, mutation group or patient ID B. Violin plots displaying the expression levels of canonical markers of representative cell types C. Distribution of cell types across the merged dataset and per sample D. UMAP visualization of all the 50,868 cells highlighting the cells annotated to the main cell types. The UMAP clusters of NEs are also marked by their most representative patient IDs

Based on canonical marker genes, we identified three major groups of cell types: neuroendocrine (NEs (markers TH, DBH, CHGB)), immune (PTPRC, CD163, CD247), and stromal (COL4A1, COL1A2) cells **(Fig1B)**. The analysis of cluster 7 revealed that it originated almost exclusively from one donor (P370) and was hallmarked by elevated expression of typical adrenocortical rather than adrenomedullar marker genes, such as CYP11A1 and CYP11B1 **(Fig. 1B-C, Fig. S1B-C)**. Hence, cells from donor P370 were considered non-representative and excluded from downstream analysis. Neuroendocrine cells (NEs) represented the largest cell fraction (63%, Clusters 0, 1, 2, 3, 4, 6, 10, 11, 16, 19), followed by the stromal (16%, Clusters 9, 12, 13, 15, 17, 18) and the immune cells (16%, Clusters 5, 8, 14, 16) (**Fig. 1C)**. Most NE clusters consisted largely of cells from single patients (**Fig. S1A-C)**. Cells of the tumor microenvironment (TME), however, occupied shared UMAP territories (**Fig. 1D)**. Based on these observations we decided not to apply batch correction in subsequent analyses to maintain the biological heterogeneity.

To obtain a more detailed insight into the cellular complexity of the TME, the immune and stromal cells were sub-selected separately for deeper analyses. Annotation of the immune cells (**Fig. S2A**) resulted in the assignment of macrophages being the major component of the immune TME [9] (expressing CD163, CDSF1R, TGFBI), followed by T-cells (CD247, IL7R, TCF7) and B-cells (MS4A1, BLK, BANK1) (**Fig. S2B**). The macrophages were the most heterogenous immune cells which could be related to tissue-specific transcriptional programs as macrophages are widely known to exert context specific functions [10, 11] (**Fig. S2A**, *arrows*). However, the adrenal macrophages (colored green) that are derived from the same location but from different tumor samples are very different. (**Fig. S2A,** *blue and red arrows).* This suggests that the macrophage transcriptome not only has a strong locational but also a tumor type-specific component. The T- and B-cells originating from different locations and mutation groups appear rather similar as they clustered together. Finally, annotation of the stromal group (**Fig. S2C-D**) revealed Schwann cells (expressing SOX6, CDH19, NRXN1), endothelial cells (FLT1, PECAM1, PTPRB) and fibroblasts (TAGLN, ACTA2, COL1A1).

The numbers of individual immune and stromal cell populations were deemed too small for in-depth analysis and were not further investigated.

### RET and SDHB Tumor Cells Display Chromosomal Aberrations

We explored inferred Copy Number Variation (iCNV) to determine large-scale somatic chromosomal changes (**Fig. 2A**). Immune- and stromal cells served as ‘reference’ in the assumption that large CNVs do not occur in the non-malignant. In agreement with published whole-genome sequencing profiles of PCPG tissue [12–15], segmental loss in the p-arm of chromosome 1 (1p) was present in all examined tumors regardless of the mutation type. Loss of 1p was not found in the TME cells confirming the assignment of the neuroendocrine cells as PCPG tumor cells. In addition, we observed widespread loss in other chromosomes for example the 3q and 6p arms as well as patient-specific aberrations such as loss in chr21 and gain in 1q, 3q, 13q and 14q (**Fig. 2A**). Apart from a few exceptions (RET-PCPG P66) we found different iCNV patterns in chr13 and chr15 in a subset of the tumor cells; in P227 (SDHB-PCPG) we identified small variations in chr3 and chr17 but observed few intra-individual heterogeneities.

**Figure2.**
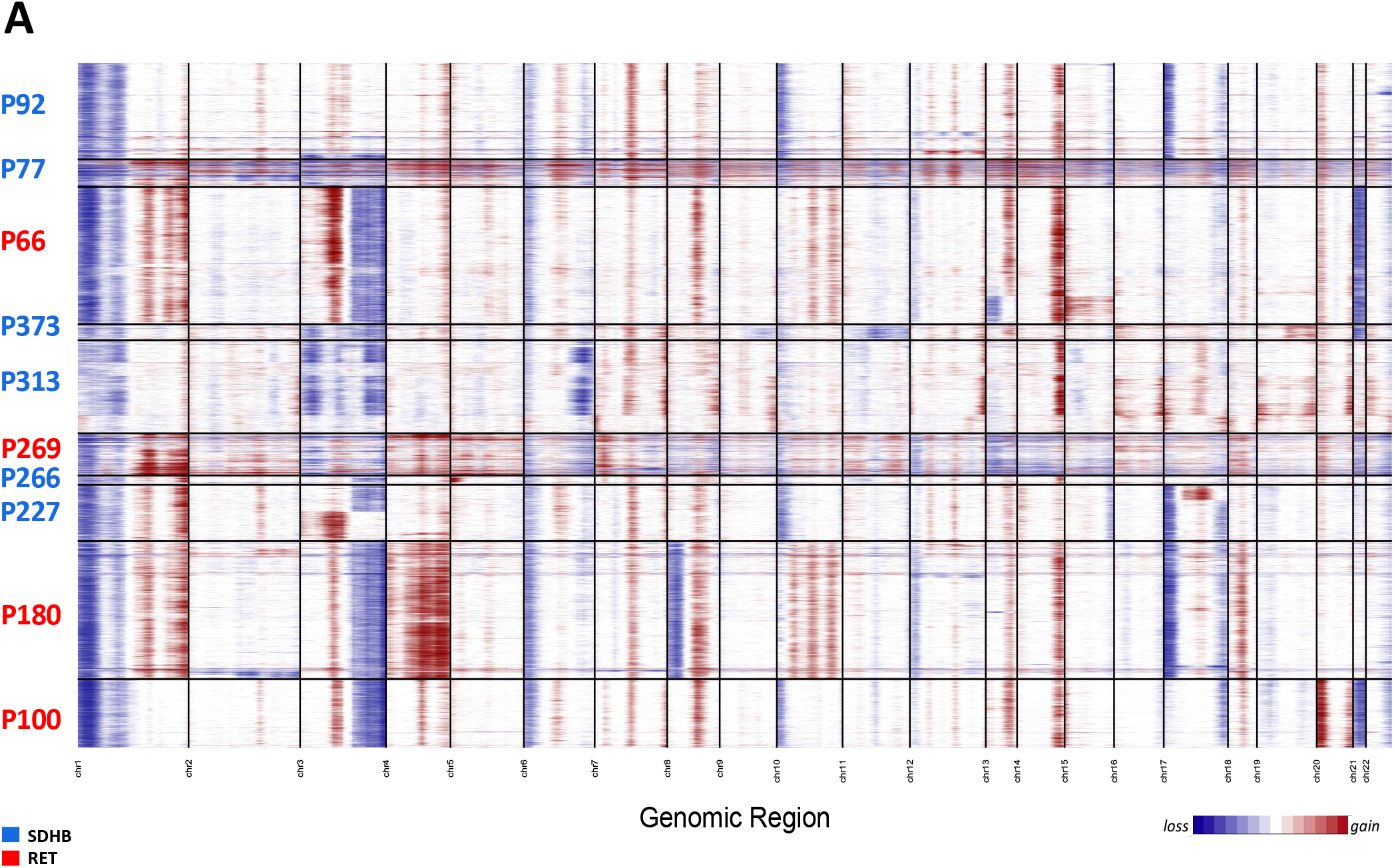
A. Heatmap of inferred CNVs of NE cells (immune clusters and stromal clusters were applied as reference). The patient IDs are colored by the mutation groups.

In contrast to the extensive inter-individual and tumor-specific genomic aberrtRET and SDHB Tumor Cellsaions, the inferred genomic profiles of tumor cells within each tumor population were largely homogeneous, suggesting that the genome remained largely stable following an initial catastrophic event, the genome remained largely stable.

### Transcription Programs Separate RET- and SDHB PCPG Tumor Cells

To assess the inter-tumoral heterogeneity between RET and SDHB PCPG tumor cells, we selected and re-clustered the tumor cells. With this finer grained resolution, we identified UMAP-clusters that consisted of cells mostly from one patient. This impinged on both the UMAP-plots annotated by patient IDs (**Fig. 3A)**and the heatmap annotation of the hierarchical clustering of top20 cluster markers (**Fig. 3B, Table S3)**, reinforcing the strong inter-individual heterogeneity observed in the iCNV analysis. Selecting the tumor cells gave us the opportunity to determine the genes that are differentially expressed between the mutation groups (**Fig. 3C**, **Table S4**). The newly identified markers were associated with either overlapping KEGG pathways (‘nervous system development’) or with gene ontology terms related to the secretory function of chromaffin cells (‘ion channel activities’, data not shown)[16].

**Figure3.**
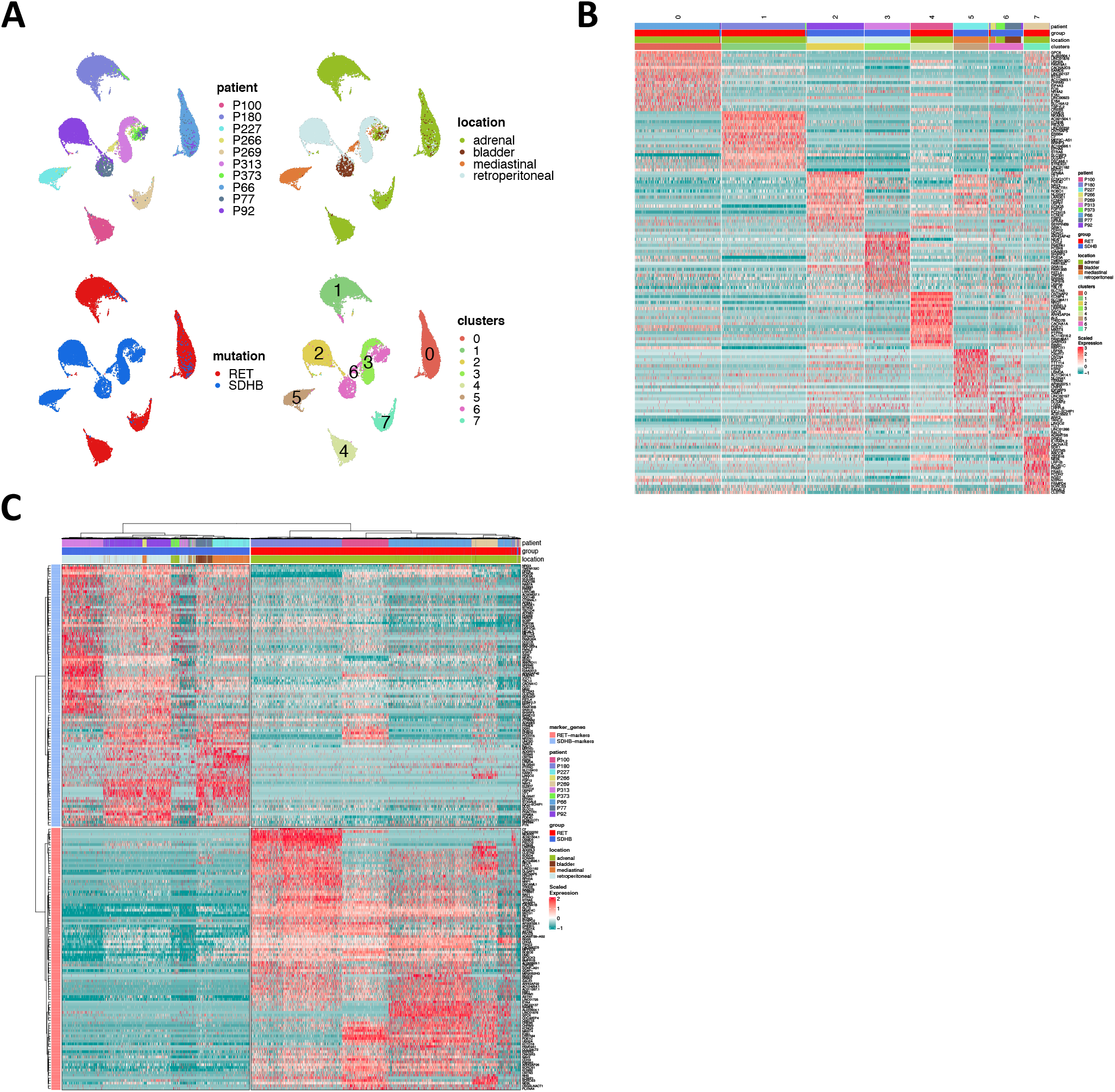
A. UMAP visualization of the PCPG tumor cells subcluster after re-clustering (no batch-correction), annotated by patient ID, tumor location, mutation group and cluster B. Hierarchical clustering of the differentially expressed genes for UMAP-clusters across the PCPG tumor cell subclusters C. Hierarchical clustering of the differentially expressed genes for RET and SDHB mutation groups (sn-markers) across the PCPG tumor cells

We wished to determine the transcriptional programs that are active across the tumor cells and then identify the programs that are differentially enriched between RET and SDHB tumors. We applied non-negative matrix factorization (NMF)[7] over the sub-selected tumor cells to determine the full transcriptional spectrum behind the intratumoral heterogeneity and to extract the most representative biological processes in the tumor cells. Firstly, we identified 28 active transcriptional programs in PCPG tumor cells of each sample, based on their transcriptional profiles at the single cell level. The signature enrichment of these 280 programs was calculated in every individual tumor cell of the whole dataset. Next, based on the enrichment scores, we hierarchically clustered the programs and identified 10 metaprograms (**Fig. 4A**). Genes were ranked according to their frequency of being present within one metaprogram. The metaprograms spanned a narrow range of functions (**Table S5**) including neuronal development (M1: BMPR1B, ROBO1; M2: NRG1, NTNG1; M3: FGF14, ROBO1; M8: SYT1, CTNNA2; M10: HDAC9, RORA), ion channel activity (M4: RYR2, PDE4B; M5: CACNA2D3, CHRM3), hormone synthesis (M9: TH, GCH1) and proliferation (M6: BRIP1, HELLS). Metaprogram seven (M7) could not be associated with a significant ontology term.

**Figure4.**
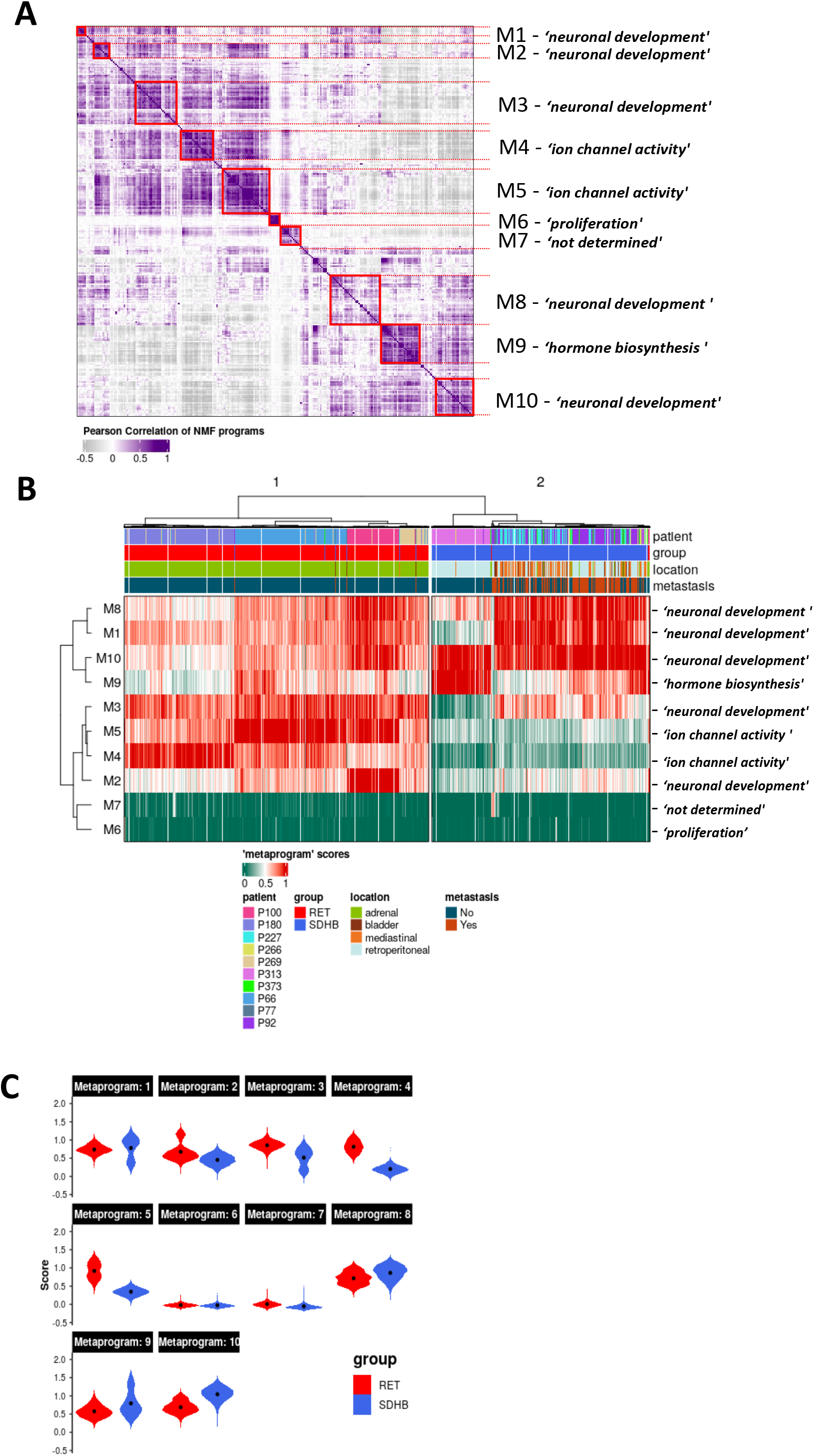
A. Heatmap showing the correlation and hierarchical clustering of the 280 factors calculated in our NMF-analysis of the tumor cells individual samples, across all mutation groups. Metaprograms are numbered M1-M10 and annotated by their representative ontology terms. B. Heatmap showing scores of PCPG tumor cells for the 10 metaprograms identified from the NMF analysis of individual samples. C. Violin-plots showing scores of PCPG tumor cells for the 10 metaprograms identified from the NMF analysis grouped per mutation group (black dots mark mean, *Wilcoxon p<2.2e-16* within each Metaprogram).

The hierarchical clustering of the metaprogram-scored cells unveiled two major clusters separating RET from SDHB tumor cells (**Fig. 4B**). The subclusters within the RET-branch segregated along the patient samples. In the SDHB-branch, however, solely sample P313 formed a discrete subcluster, while the tumor cells of other SDHB patients formed mixed subclusters. Surprisingly, the SDHB tumor cells (originating from various anatomical locations) are less heterogenous than their RET counterparts (originating from the adrenal gland).

The most pronounced differences in the average enrichment scores between the RET (cluster 1) and the SDHB (cluster 2) cluster were evident at metaprograms M2, M3, M4 and M5 (**Fig. 4C**). The ‘ion channel activity’ of the M4-M5 metaprograms is highly enriched among the RET tumor cells indicating a high secretory activity of the adrenal RET-pheochromocytoma tumor cells. The M9 ‘hormone synthesis’ program was more enriched among the SDHB tumor cells, but mainly due to patient P313. A very small fraction of cells scored high for the ‘proliferation’ (M6) metaprogram, revealing a low but appreciable proliferative capacity of the PCPG tumor cells. Several metaprograms were associated with ‘neuronal development’ ontology terms and were shared in both branches of the tumor group separations.

In sum, the NMF analysis revealed two main transcriptional programs in PCPG that separated RET from SDHB tumor cells. Genes associated with ion channel activities (secretion) were more enriched in RET tumors. We also observed that ‘neuronal development’ was a highly represented transcriptional program in both PCPG tumor cells.

### PCPG Tumor Cells Display Early Adrenal Developmental Signatures

The NMF analysis revealed several metaprograms that were associated with neuronal development but showed different enrichments scores among the mutation groups. This implies the developmental signature as an important element of the tumor cells’ transcriptome, but the differences between the mutation groups were not reflected in the ontology terms. To shed light on the developmental aspects of the SDHB and RET-PCPG tumors, we compared the transcriptome of the cell types identified in the developing human adrenal gland (8-21 weeks [17] with the PCPG tumors. We applied logistic regression and calculated probability scores for cell type matches (**Fig. 5A**). The analysis revealed that tumor cells were most similar to the cells at the junction to sympathoblast and chromaffin cells, called the ‘bridge cells’ [18]. The cell types of the PCPG microenvironment showed high similarity with their normal cell counterparts in the developmental adrenal gland dataset.

**Figure5.**
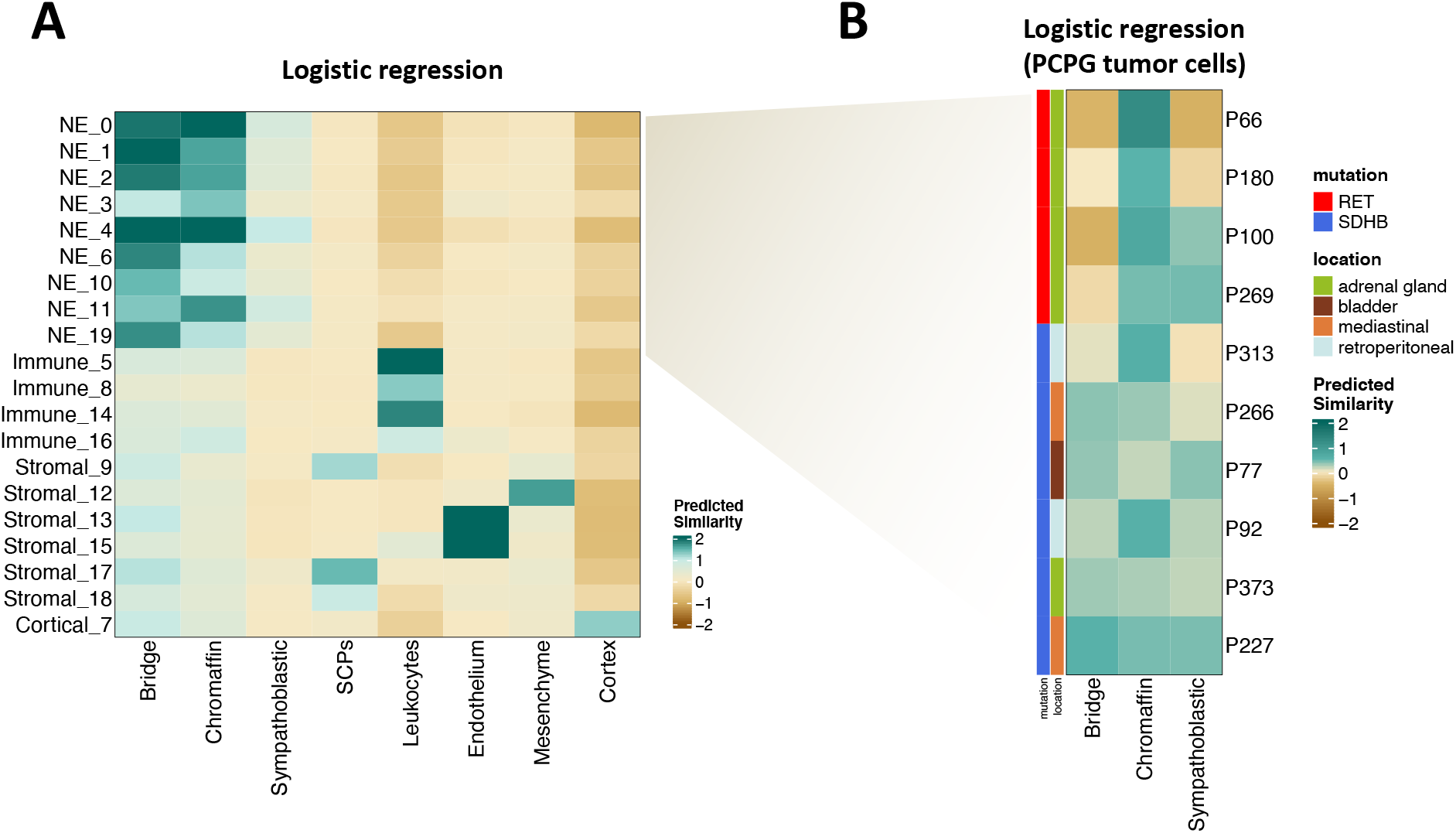
A. Heatmap showing similarity scores (logistic regression and logit scale) of the signatures of developing cell types from [17] (fetal adrenal dataset) (x axis) to PCPG cells (y axis) B. Heatmap showing similarity scores (logistic regression and logit scale) of the signatures of developing adrenal cell types from [17] (fetal adrenal dataset) (x axis) to PCPG tumor cells, by patient (y axis)

The difference between the SDHB and RET-PCPG became even more evident when the tumor cells of each patient were compared to the chromaffin developmental cell types (**Fig. 5B**). The logistic regression confirmed that the RET-PCPGs were more similar to the reference chromaffin cells, while the SDHB-PCPGs scored highest with both the chromaffin and the bridge cell types, suggesting an earlier developmental state.

## DISCUSSION

We performed snRNA-seq and mapped the transcriptional landscape of PCPG to investigate the tumor heterogeneity and to identify the transcriptomic programs that are associated with the mutation group of the tumor. We explored the transcriptional heterogeneity by the analysis of the transcriptomic profiles of 50,868 single nuclei from 11 patients (counting all cell types from 5 RET- and 6 SDHB PCPG tissue samples). This is the first study unveiling the PCPG heterogeneity and the consequences of germline mutations at the single-cell level.

Neuroendocrine cells, the largest population in the dataset, were identified as tumor cells on the basis of marker genes and in particular by inferring copy number variations from gene expression levels [19]. The iCNV profiles revealed two important features: firstly, the lack of tumor cell sub-clusters within patients suggests a single initial catastrophic event that led to the birth of the tumor cells. Secondly, apart from very few recurring aberrations, we identified rather patient-specific iCNV patterns, marking the level of inter- and intratumoral heterogeneity in PCPG cellulome that provided a challenge for tumor classification.

To identify the patterns of the single-nuclei transcriptomic profiles based on the tumor cells, we applied NMF, an unsupervised learning approach that is employed to approximate high-dimensional datasets in a reduced number of meaningful components [7, 20, 21]. The analysis of single nuclei transcriptomes of >30,000 tumor cells resulted in 10 metaprograms across the entire tumor set. The transcription programs related to ion channel activity (transmembrane transport) separated SDHB and RET tumor cells. Based on biochemical analysis of plasma, urinary and tissue samples, we previously [22] found that RET tumors produce (and contain) higher concentrations of catecholamines but secrete them at a lower rate than SDHB tumors. Our cohort was not split by the hormone synthesis metaprogram, moreover (due to a single patient) it showed a higher mean enrichment in the SDHB subset. However, it was split along the ion channel (transmembrane transport) programs that we associated with secretion [23]. Metaprograms linked to neuronal development were found to be active throughout the tumor cells irrespective of their mutational groups.

To explore the developmental status of the tumor, we utilized published datasets of the developing adrenal gland as reference. Applying logistic regression revealed that the RET-PCPG tumor cells are transcriptionally more similar to developed adrenal chromaffins, while SDHB-PCPG tumor cells appear to be in an earlier phase of adrenal development. Our results suggest that PCPG tumor cells had a primarily chromaffin-like phenotype suggesting that the chromaffin cell development state maybe related to the mutation-associated prognosis.

In summary, we revealed extensive levels of heterogeneity among PCPG tumor cells and identified transcriptional programs related to neuronal development as key processes active in these tumor cells. We speculate that in RET-PCPG, the mutation caused a development block during late chromaffin development as compared to the ‘more immature’ SDHB-PCPG tumors. To differentiate this developmental block from alternative transformative events that could also lead to the modified transcriptomes of the tumor cells, investigation of larger cohorts is needed. Understanding the origin of the tumor and the sources of its heterogeneity may help the development of targeted therapies.

## DATA AND MATERIAL’S AVAILABILITY

### Lead Contact

Further information and requests for resources and reagents should be directed to and will be fulfilled by the Corresponding author.

### Materials Availability

This study did not generate new unique reagents.

### Data and Code Availability

The high-throughput datasets have been deposited in the European Genome-phenome Archive. The accession number for the single cell expression data of PCPG tumor samples reported in this study is EGAS00001006230. This study did not generate any unique code. All software tools used in this study are freely or commercially available.

## ACKNOWLEDGEMENTS

The authors would like to thank the patients, their families for supporting this research. Our thanks are extended to Igor Adameyko *(Karolinska Institute, Stockholm, Sweden)* for discussions.

## ETHICS DECLARATIONS

### Ethical Statement

The study was approved by the Institutional Review Board of the Radboud University Medical Center and informed consent was obtained from each patient (protocol no 9803-0060)

**Human Tumor Samples**

Tumor tissue samples of patients with a biochemically / histologically proven diagnosis of pheochromocytoma/paraganglioma were investigated. Fresh samples were snap frozen, usually within 1 hour after resection.

---

**FigureS1.**
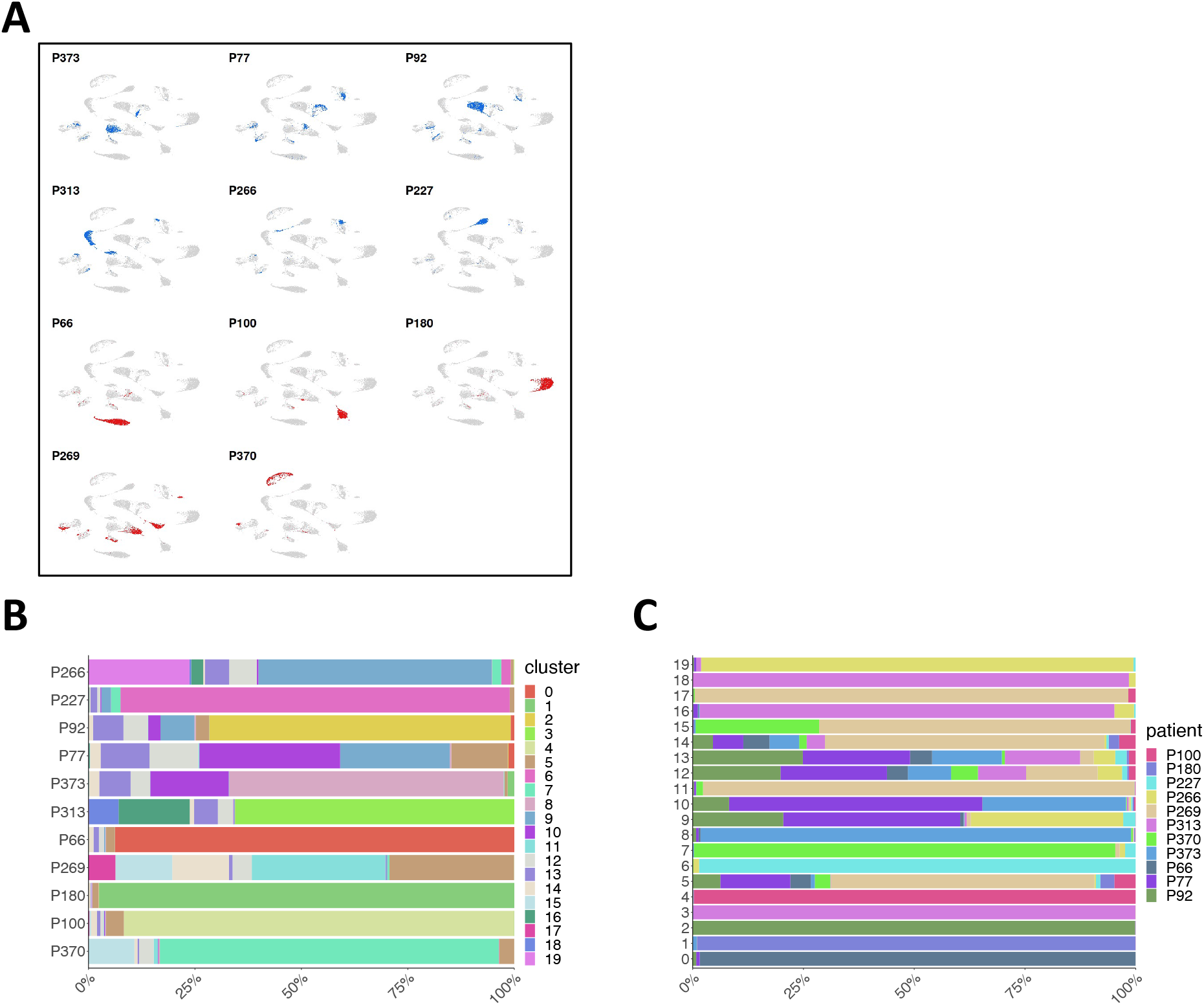
A. UMAP visualization of the merged dataset separately annotated by patient (blue: ‘SDHB group’, red: ‘RET-group’) B. Fraction of cells per sample populating the UMAP-clusters C. Fraction of cells per UMAP-clusters found per sample

**FigureS2.**
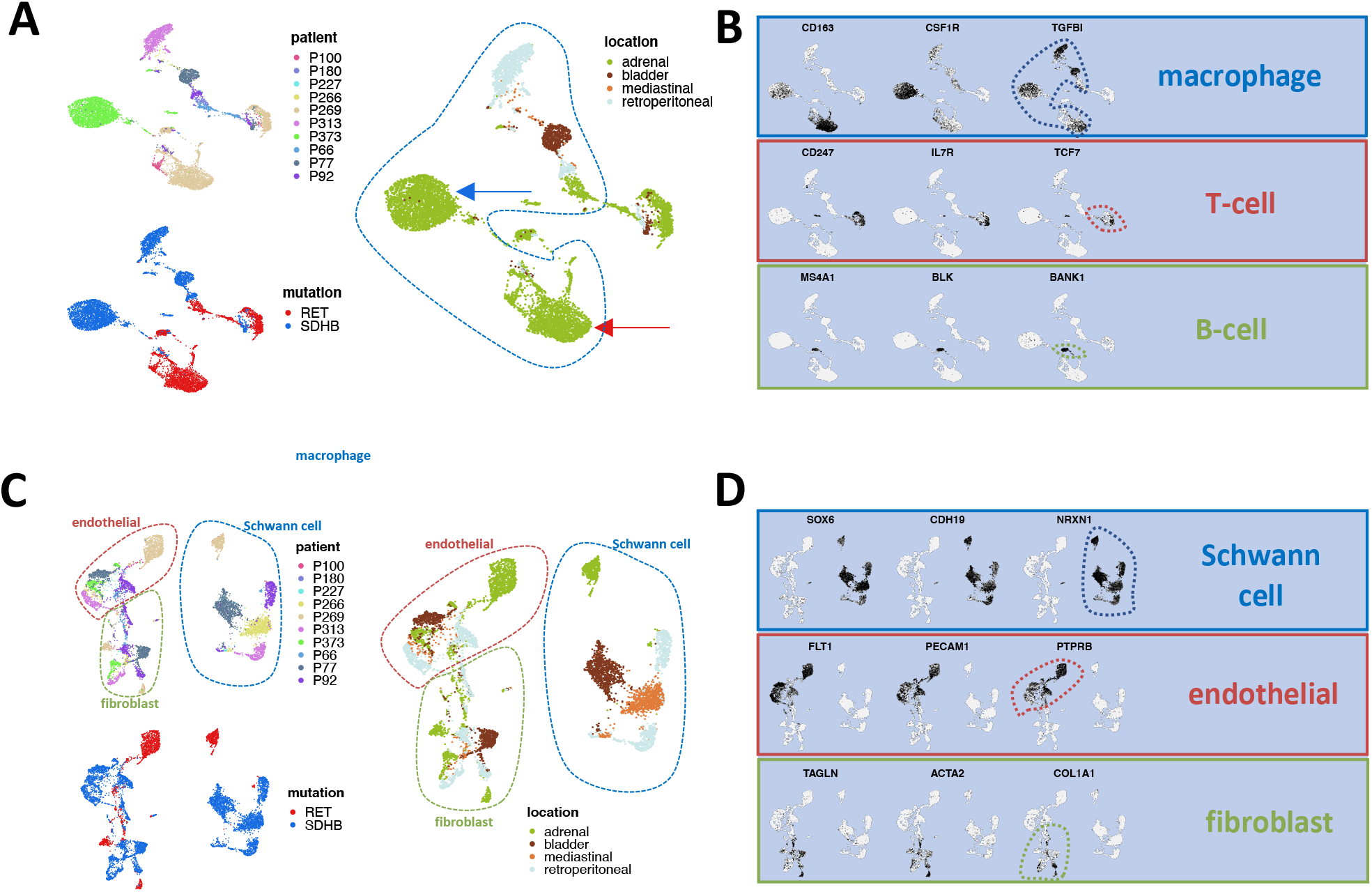
A. UMAP visualization of the PCPG immune cells subcluster after re-clustering (no batch-correction). The arrows point at the cells annotated as macrophages, found in tumors from similar anatomical locations. (blue: from an SDHB-tumor, red: from a RET-tumor) B. UMAP visualization and the relative expression levels of canonical cell type markers across the PCPG immune cells subcluster C. UMAP visualization of the PCPG stromal cells subcluster after re-clustering (no batch-correction) D. UMAP visualization and the relative expression levels of canonical cell type markers across the PCPG stromal cells subcluster

## Additional Data

**Table S1**. Clinical information and snRNAseq quality parameters of processed/analyzed samples

**Table S2**. Top50 differentially expressed markers of the 20 clusters in the complete merged dataset

**Table S3**. Top50 differentially expressed markers of the tumor cell sub-clusters **Table S4**. Differentially expressed markers of the of mutation groups based on the tumor cells

**Table S5**. Gene lists of the 10 metaprograms identified in the tumor cells. The cells with unique markers (across the metaprograms) are colored blue.

